# A State-Dependent Drift Diffusion Model Reveals Mice Actively Explore A Speed–Accuracy Continuum During Decision-Making

**DOI:** 10.1101/2025.06.02.657156

**Authors:** Ryan A. Senne, Cristina Delgado-Sallent, Steve Ramirez, Benjamin B. Scott, Brian DePasquale

## Abstract

Understanding how animals shift between different decision-making strategies is critical for bridging normative models with naturalistic behavior. While drift diffusion models (DDMs) provide a powerful framework for describing evidence accumulation in two-alternative forced choice (TAFC) tasks, they assume fixed parameters across trials—an assumption often violated in practice. Here, we introduce a state-dependent DDM framework in which discrete latent states modulate decision parameters from trial to trial. This approach reveals that mice dynamically switch between impulsive and deliberative decision states that differ in accuracy and response latency, suggesting active exploration of the speed–accuracy trade-off. We uncover rare high-bound states in which mice exhibit deliberation times and accuracies approaching those observed in humans. These results raise new questions about the cognitive flexibility of rodent decision-making and offer a foundation for studying how internal states and external variables—such as reward history or uncertainty—influence strategy selection. Our method provides a natural interface for integration with neural recordings and dynamical systems models, offering a path toward identifying the circuit-level mechanisms underlying adaptive decision behavior.

## 1 Introduction

A qualitative description of behavior is a major goal of neuroscience [1], psychology [2], and economics [3]. Historically, there has been a focus on normative behavioral models, but with greater tools for behavioral tracking [4] and larger behavioral datasets [5], there has been significant progress in the development and application of descriptive models of behavior [6, 7]. When applied to decision-making behavior these approaches have identified several behavioral strategies employed by many species, including humans, rats and mice, that deviate from normative behavior, including stochastic [6] and outcome-dependent [8] trial sequence dependent shifts in strategy. The success of these approaches supports further development of similar approaches.

A cornerstone of normative models of decision making is the drift diffusion model (DDM) [9, 10], where the cognitive process that underlies a decision is modeled as a noisy accumulation to a bound. This model is optimal under several assumptions, such as stationarity, [11], and as a result it has been widely influential in the field of cognitive decision making and psychology. However, recent evidence suggests that many animals deviate from normative assumptions while making decisions [12].

Here we present a state-dependent modeling framework to account for trial-history dependent shifts in decision making that otherwise conform to a normative decision-making strategy defined by the DDM. To do so, we augment the DDM to include a underlying discrete latent process that changes the DDM parameters from trial to trial.

Using our method we identified that mice performing a two alternative forced choice task shift decision making strategy from trial to trial. Our model improves reaction time distribution prediction compared to a classical DDM and provides insight into the shifting decision making strategy of animals while maintaining the desirable normative quality of the DDM. Using our framework we discovered that mice exhibit different behavioral states with dramatically different reaction time distributions and and accuracies, suggesting active exploration of the speed–accuracy trade-off. Interestingly, our method finds rare states with high decision bounds, suggesting that mice may alternate between different decision policies as they perform operant tasks.

These results suggest that even simple perceptual decision-making tasks engage a repertoire of internal strategies that fluctuate over time, reflecting dynamic shifts in cognitive state. By capturing these latent dynamics, our framework offers a window into how animals balance competing goals such as speed and accuracy, and highlights the potential for richer structure in behavior than is typically modeled. More broadly, this approach opens the door to identifying state-dependent computations in neural activity and provides a powerful tool for linking behavior to brain-wide decision circuits.

## 2 State-dependent drift diffusion model

We consider a hierarchical time-dependent probabilistic model to describe the decisions and reaction times of a subject performing a binary decision-making task (Figure 1 A). For a sequence of behavioral trials, we seek to identify a latent state sequence 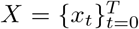 that defines the subject strategy, given a set of observations 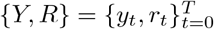, where *y*_*t*_ ∈ R_+_ is the response time and *r*_*t*_ ∈ {− 1, +1} is the observed choice respectively on trial *t*. We assume the value of the latent state on trial *t x*_*t*_ is conditionally dependent on the previous value, *x*_*t*_ *−*_1_. Below we detail how we model the latent state sequence and how this relates to the set of observations.

**Figure 1.**
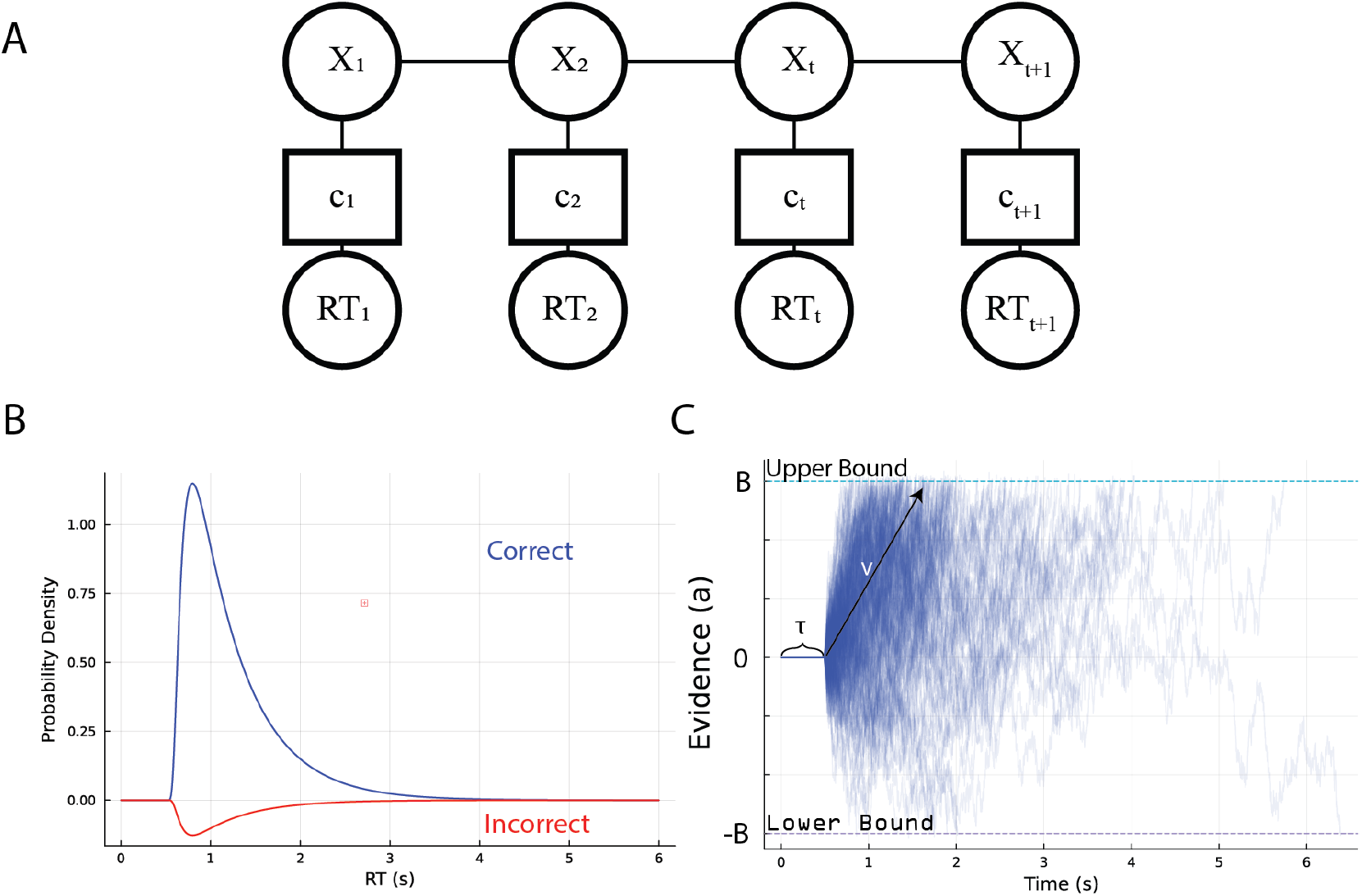
State-dependent DDM architecture. **A.** Factor graph of the state-dependent DDM model. Reaction times (RT) and choice (c) are conditionally dependent through the latent variable *X*. **B**. Wiener First Passage Time (WPFT) distribution. Analytical solution of the reaction times for the canonical DDM model. **C**. Simulations of a single-state DDM process. Evidence starts at some initial value, and does not accumulate until surpassing some non-decision time, *τ*. After this, evidence accumulates with drift *v* noisily until reaching the symmetric absorbing bounds at *B* or −*B*.

### 2.1 Between trial discrete latent dynamics

Following recent work that suggests animals and humans employ a discrete set of behavioral strategies during decision making tasks [6] and given our assumption of Markovian temporal dynamics, we model the latent state dynamics using a hidden Markov model (HMM). A *K* state HMM is specified by a set of parameters *θ* = {*π, A, ϕ*}, where *π* is the initial state distribution, *A* determines that state transitions, and *ϕ* parametrizes the observation distribution for {*Y, R*}. For every state *i* and pair of states *i, j* ∈ 1, …, *K*,

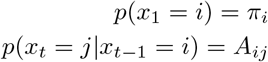

The emission distribution *p*(*y*_*t*_, *r*_*t*_ | *x*_*t*_, *ϕ*), which we specify below, is generic but assumes that *y*_*t*_ and *r*_*t*_ is conditionally independent of previous observations given *x*_*t*_.

### 2.2 Within trial evidence accumulation: Drift diffusion model

The optimal decision model for the two-alternative forced-choice (TAFC) task we consider here is the drift diffusion to bound model (DDM) [10]. In this model, a latent decision variable *a*(*t*) of accumulated evidence evolves over time as noisy, biased evidence accumulates (Figure 1 C). The evolution of this process is governed by the following stochastic differential equation:

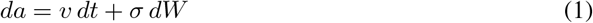

where *v* is the average rate of evidence accumulation, called the drift rate, *σ* is the diffusion coefficient capturing the noise magnitude, and *dW* is an increment of a Wiener process (i.e., Brownian motion). Evidence accumulation continues until *a*(*t*) reaches one of two absorbing boundaries, set symmetrically at ± *B*. Crossing the upper boundary corresponds to a correct response, while crossing the lower boundary corresponds to an incorrect response. This setup makes the DDM particularly suitable for TAFC tasks, as it models both the binary decision outcome (e.g., correct/incorrect) and the continuous distribution of response times (RTs; Figure 1 B), capturing both accuracy and latency.

We include two additional parameters in the model, consistent with prior implementations of the DDM. *τ* (Figure 1 C) indicates the time between stimulus onset and evidence accumulation, referred to as the ‘non-decision time’. *a*_1_ ∈ [0, 1] denotes the relative starting point between the two boundaries.

For every state *i* of our latent dynamics model, we define a individual DDM observation model with a unique set of parameters. We aggregate the DDM parameters for state *i* into a set of observation parameters *ϕ*_*i*_ = {*v*_*i*_, *B*_*i*_, *σ*_*i*_, *τ*_*i*_, *a*_1*i*_}.

## 3 Inference and learning

We define all model parameters as 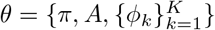. We refer to the above formulation of our model as a HMM-DDM, given its unique combination of a HMM with a DDM observation model. We are primarily interested in using our model to identify parameters that best describe our data, and to infer the most likely latent state sequence *X*. Direct maximum likelihood estimation is intractable given our model’s graphical structure. Instead we use the well-known expectation-maximization (EM) algorithm.

In EM, we do not maximize the log-likelihood function directly but instead maximize an auxiliary function, referred to as the Q-function, defined as:

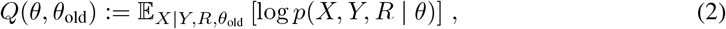

where 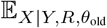 is the expectation over the latent sequence *X*. Given the factor graph structure of our model (Figure 1 A), we write the log joint likelihood as

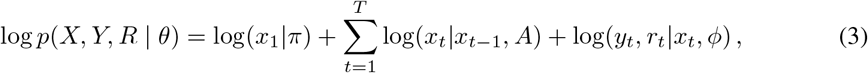

The DDM log likelihood for trial *t* log *p*(*y*_*t*_, *r*_*t*_ | *ϕ*) is calculated using closed-form expressions from the Wiener first-passage time (WFPT) distribution (Figure 1 B) which is derived analytically via solutions to the Fokker-Planck equation [13]. Although the exact solution involves an infinite series, fast and accurate numerical approximations are available, making this computationally tractable [10, 13].

### Inference and M-step

To infer the latent state trajectory *X* we use the standard forward-backward algorithm for HMMs [14]. During the M-step *π* and *A* are updated using closed form updates for HMMs [14]. We update each state’s DDM parameters *ϕ*_*k*_ by maximizing its responsibility-weighted log-likelihood with L-BFGS:

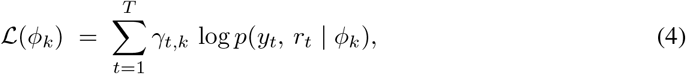

where *γ*_*t,k*_ = *p*(*x*_*t*_ = *k* |*Y, R*) is the posterior responsibility obtained from the forward–backward pass. For all states, we do not optimize *σ* or *a*_1_ due to non-identifiability; instead, we fix *σ* = 1.0 and *a*_1_ = 0.5

## 4 Model Implementation

All models and analyses were implemented in the Julia programming language[15]. The complete source code and reproducibility materials are publicly available at: https://github.com/rsenne/ DriftDiffusionModels.jl. The HMM backend was implemented via HiddenMarkovModels.jl [16].

## 5 Related work

A number of studies have developed and applied a variety of hierarchical decision-making models to data from animals and humans. The Python-based toolbox ‘HDDM’ (hierarchical drift diffusion model) [17, 18] uses a hierarchical Bayesian framework that pools information across participants, simultaneously estimating group-level and individual drift-diffusion parameters, providing increased statistical power actually. Because each person’s parameters borrow strength from the shared group priors, the toolbox can fit large multi-subject datasets—even when each participant contributes only a modest number of trials—while returning full posterior uncertainty for every level of the hierarchy. Because HDDM fixes each subject’s latent DDM parameters and assumes trials are conditionally independent given those constants, its hierarchical graph lacks the state-space layer we describe here required to capture time-evolving dynamics within a single subject.

The approach of [8] shows that choice biases and seemingly stimulus-independent errors (‘lapses’) can be explained by a between-trial latent process: an exponentially filtered memory of previous choice-outcome pairs that defines the initial state of the DDM for each trial. However, because the latent variable modulates only the accumulator’s starting point while keeping the decision bound fixed, the approach cannot account for trial-wise adaptations that act by raising, collapsing, or otherwise adjusting boundary height, as we consider here. Other studies have identified trial-to-trial fluctuations in the bound height can account for changes in decision-making strategy[12], suggesting the more flexible DDM observation model of our approach is necessary to fully capture the statistics of decision-making across trials.

Finally, [6] fit mouse choice sequences with a similarly structured latent variable model, except that a Bernoulli generalized linear model (GLM) with various regressors, such as stimulus strength and reward history, was used as the observation model. The proposed ‘GLM-HMM’ model also infers the latent ‘strategy’ state path using a hidden Markov model, and learning is done via EM. However, because the GLM-HMM outputs only binary choices and lacks an explicit accumulation-to-bound mechanism, it cannot generate or fit the reaction-time distributions our state-dependent DDM can, limiting its ability to explain speed-accuracy trade-offs.

## 6 Applications to synthetic data

We first bench-marked our model on synthetic data generated from a multi-state DDM. The latent state sequence was generated from a HMM with two states. We evaluated two datasets: one in which the two DDMs had similar parameters (referred to as ‘low separation’, indicating the model parameters not strongly separated), and another in which the DDMs were highly distinct (‘high separation’).

For low separation, both states corresponded to engaged behavior, i.e., DDM parameters that produce slow reactions times and high accuracy, while for high separation, one state reflected engaged behavior and the other disengaged behavior, which produced fast reaction times and low choice accuracy (Figure 2 B). For both settings, we used the same initial state distribution *π* and transition matrix *A* (Figure 2 A) to generate the data, and used 10K trials in both cases. We selected these two conditions to illustrate parameter recovery under relatively difficult and easy conditions, respectively.

**Figure 2.**
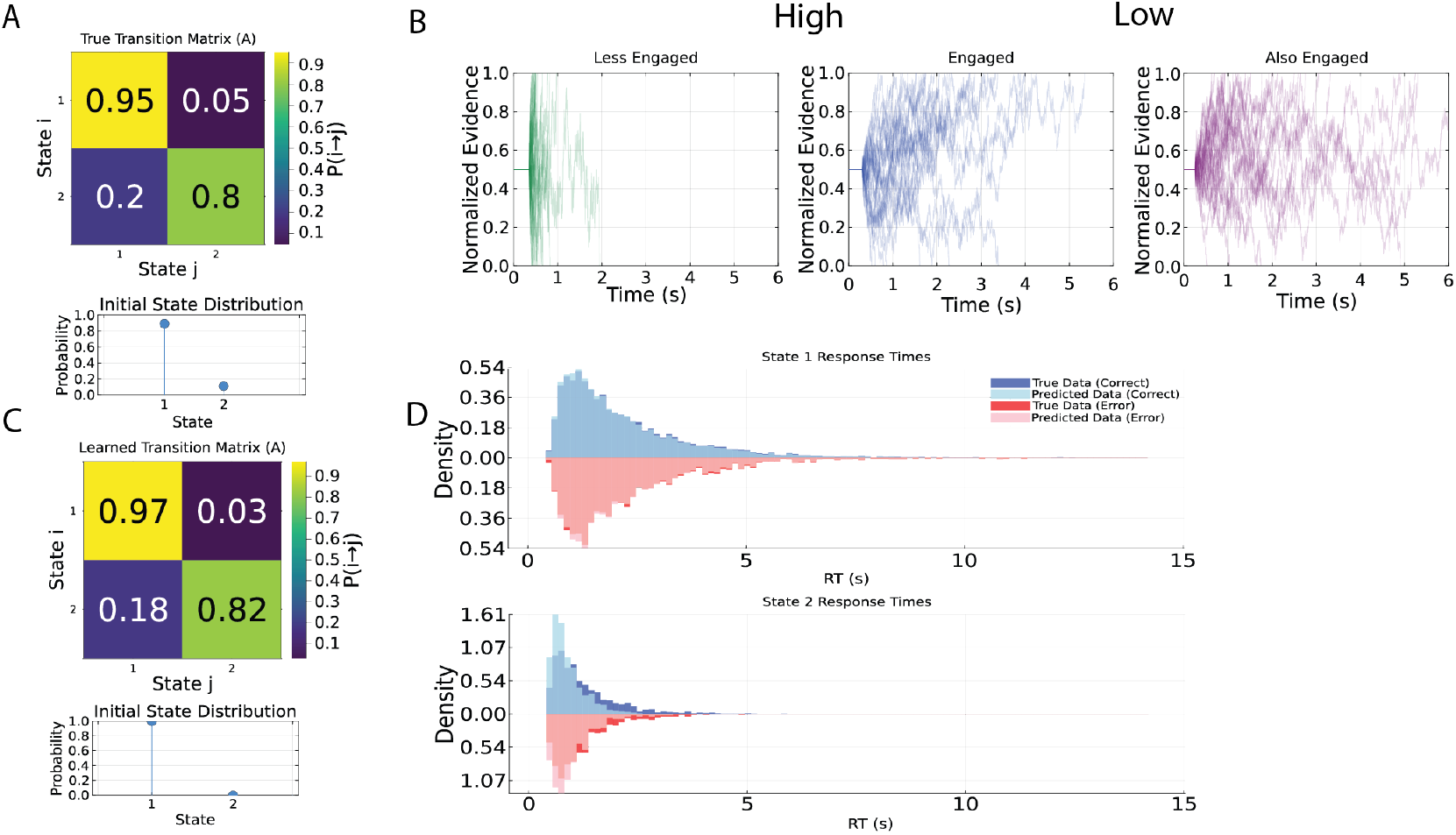
Parameter recovery of the model. **A.** True model parameters of the HMM. Top is the transition matrix *A* and bottom is the initial state *π*. **B**. Possible DDM emission models that the HMM can generate. We consider two models: a ‘high separation model’ where the two DDMs represent an engaged and disengaged strategy, respectively, or ‘low separation models’ where both DDMs produce similar reaction times distributions. **C**. The learned HMM parameters for the ‘high separation’ model. **D**. True and predicted reaction times from the learned model for the high separation synthetic data. The overlapping distributions indicate that the true and predicted RTs are very similar, and thus model recovery was achieved.

To assess the model’s ability to recover parameters under different data regimes, we varied the sequence length for values of *T* ∈ {10^1^, 10^2^, 10^3^, 10^4^, 10^5^} (Figure 3). When sufficient data were available (e.g., 10K trials), the model reliably recovered accurate point estimates of the transition matrix (Figure 2 C), effectively captured the underlying DDM parameters, and was able to predict response time distributions for both correct and error trials for all latent states (Figure 2 D).

**Figure 3.**
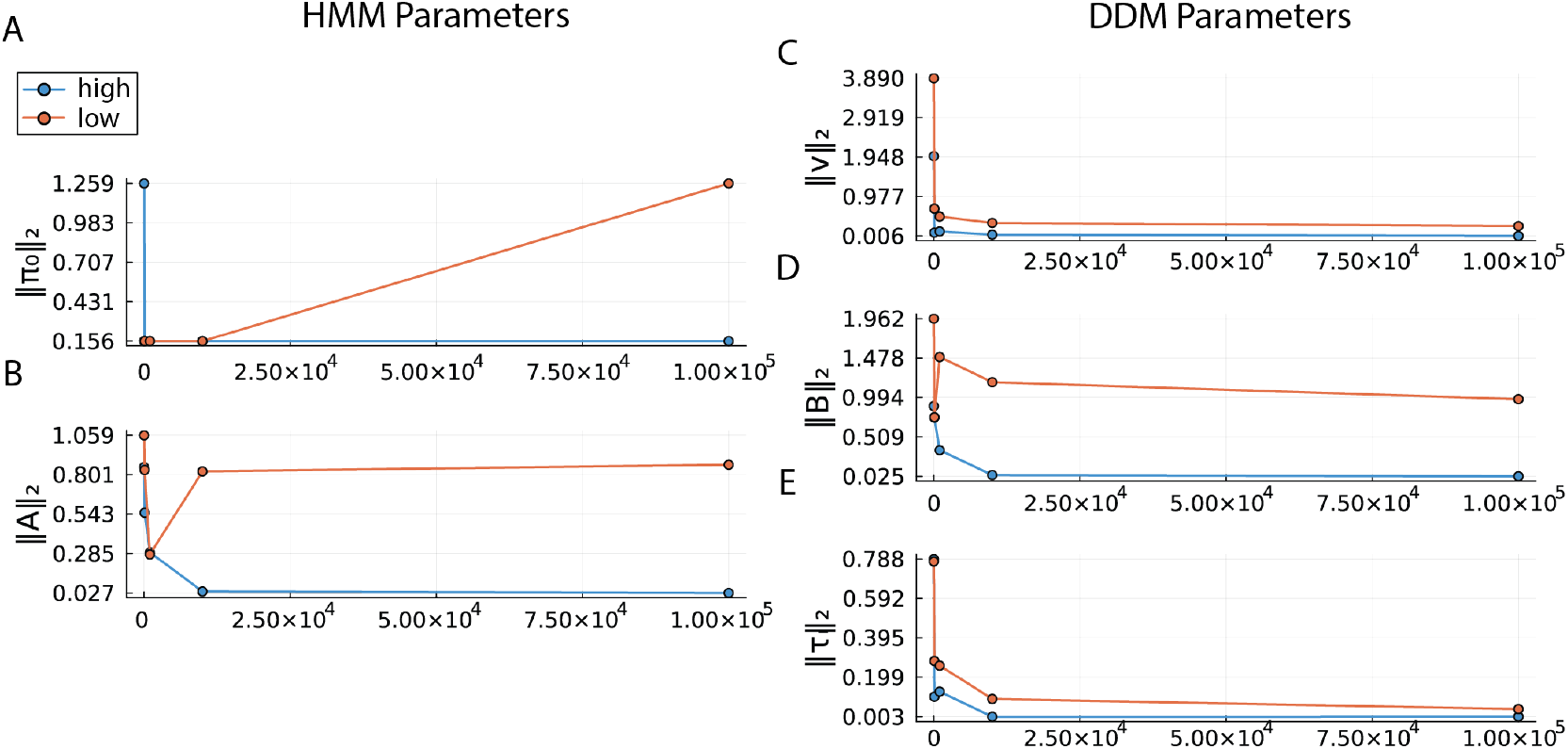
Parameter error estimation as a function of latent state sequence length. **A.** Mean squared error (MSE) of the initial state distribution *π* between the generated and recovered parameters. **B**. MSE of the transition matrix *A* between the generated and recovered parameters. **C**. MSE of the drift rates between the generated and recovered parameters. **D**. MSE of the boundary separations between the generated and recovered parameters. **E**. MSE of the non-decision times between the generated and recovered parameters.

As expected, the mean squared error of the estimated parameters decreased consistently with increasing sequence length (Figure 3). We also observed that the high separation model—where the latent states differed more markedly in behavioral signatures—yielded lower parameter estimation error than the low separation model, highlighting the benefits of state distinctiveness for model identifiability.

## 7 Application to mouse decision-making task

We next applied our method to behavioral data collected from mice performing a 2AFC task to determine if mice exhibited multiple decision-making strategies defined by different reaction time distributions. Previous approaches applied to similar tasks have identified behavioral strategy switches, but have not accounted for changes in reaction time distribution due to the model structure [6].

The data we analyzed came from three water-restricted mice performing a self-paced decision task using three nosepoke ports arranged in a horizontal row, with the center port flanked symmetrically by left and right ports spaced about 25 mm apart (Figure 4). A trial began when the animal entered the illuminated center port, triggering a sequence of 100 ms LED flashes to the left or right port drawn from an 80:20 Bernoulli process that favored a randomly chosen correct side. The flash stream ceased once the mouse selected a side port: correct choices delivered 5 µL of 10% sucrose and 3 s of light, incorrect choices triggered a 5 s lights-off timeout, and omissions were registered if no response occurred within 8 s. The details of data collections can be found in [19], and the data was provided by the authors (by request). The data are summarized in Table 1.

**Table 1:** Summary of behavioral performance across three mice.

|  | # of sessions | # of trials | Accuracy (%) | Avg. reaction time (ms) |
| --- | --- | --- | --- | --- |
| FTRAP-7 | 22 | 10 133 | 68.9 | 476 |
| MABL6-9 | 21 | 12 125 | 66.3 | 422 |
| MTRAP-2 | 22 | 9 590 | 62.2 | 403 |

**Figure 4.**
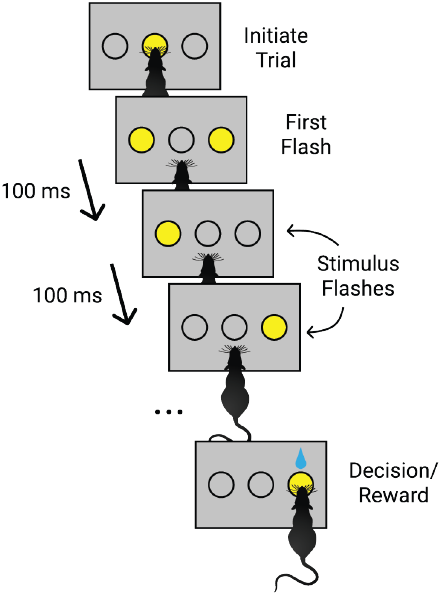
Mouse perceptual integration task. Mice initiate a trial by nose-poking a center port. Flashes occur every 100 ms in one of two bilateral flash ports. The trial ends when the mouse chooses the left or right port. Adapted from [12].

We fit our state-dependent DDM to these data to determine if mice change decision making strategy during behavioral sessions, and if these changes impacted their speed-accuracy tradeoff. To determine the appropriate number of latent states for modeling the data, we drew on previous work indicating that mice typically exhibit 2–3 distinct behavioral states during decision-making tasks [6, 19]. We fit HMM-DDMs with *K* = {1, 2, 3} latent states and evaluated model fit using the Akaike Information Criterion (AIC) and Bayesian Information Criterion (BIC). Both criteria favored models with more latent states: the two-state model outperformed the one-state model, and the three-state model further improved upon the two-state model (Figure 5 A).

**Figure 5.**
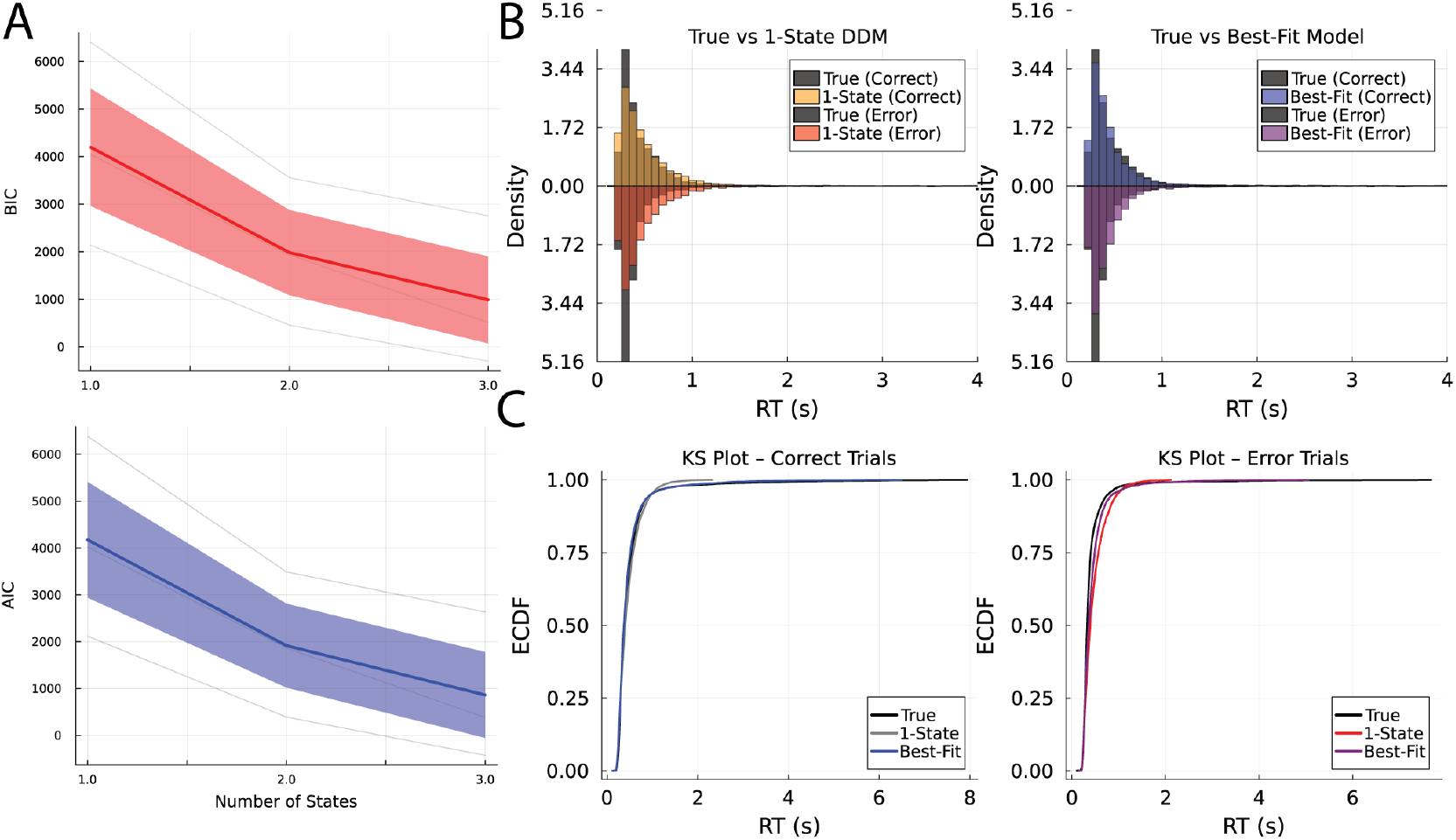
Goodness of fit metrics. **A.** Akaike and Bayesian information criteria for HMM-DDMs with 1,2, and 3 latent states. Higher state dimensions are better fit to the data. **B**. Posterior predictive checks of empirical reaction time distributions. Simulated data from the best-fitting three state model (right) better fit the empirical distributions. **C**. Kolmogorov–Smirnov (KS) plots of the empirical data from the one and three state model to the true data across error and correct trials. The three state model shows less deviance from the empirical CDF, supporting a three state model as best fit to the data.

To validate these results, we conducted a posterior predictive check. We simulated response times and choices from the fitted one- and three-state models, and compared these to the empirical reaction time distribution (Figure 5 B). The three-state model more accurately reproduced the empirical distribution, a result further supported by Kolmogorov–Smirnov (KS) plots, which showed reduced deviation between the model-generated and empirical cumulative distribution functions (Figure 5 C). These findings suggest that incorporating three latent states corresponding to three distinct behavioral strategies provides a better fit to the behavioral data.

### Characterizing the behavioral states

To interpret the behavioral significance of the latent states, we examined the response time (RT) distributions predicted by the fitted DDM for each state (Figure 6 A). The three inferred states exhibited distinct patterns consistent with different positions along a speed–accuracy tradeoff (Figure 6 B and 6 C).

**Figure 6.**
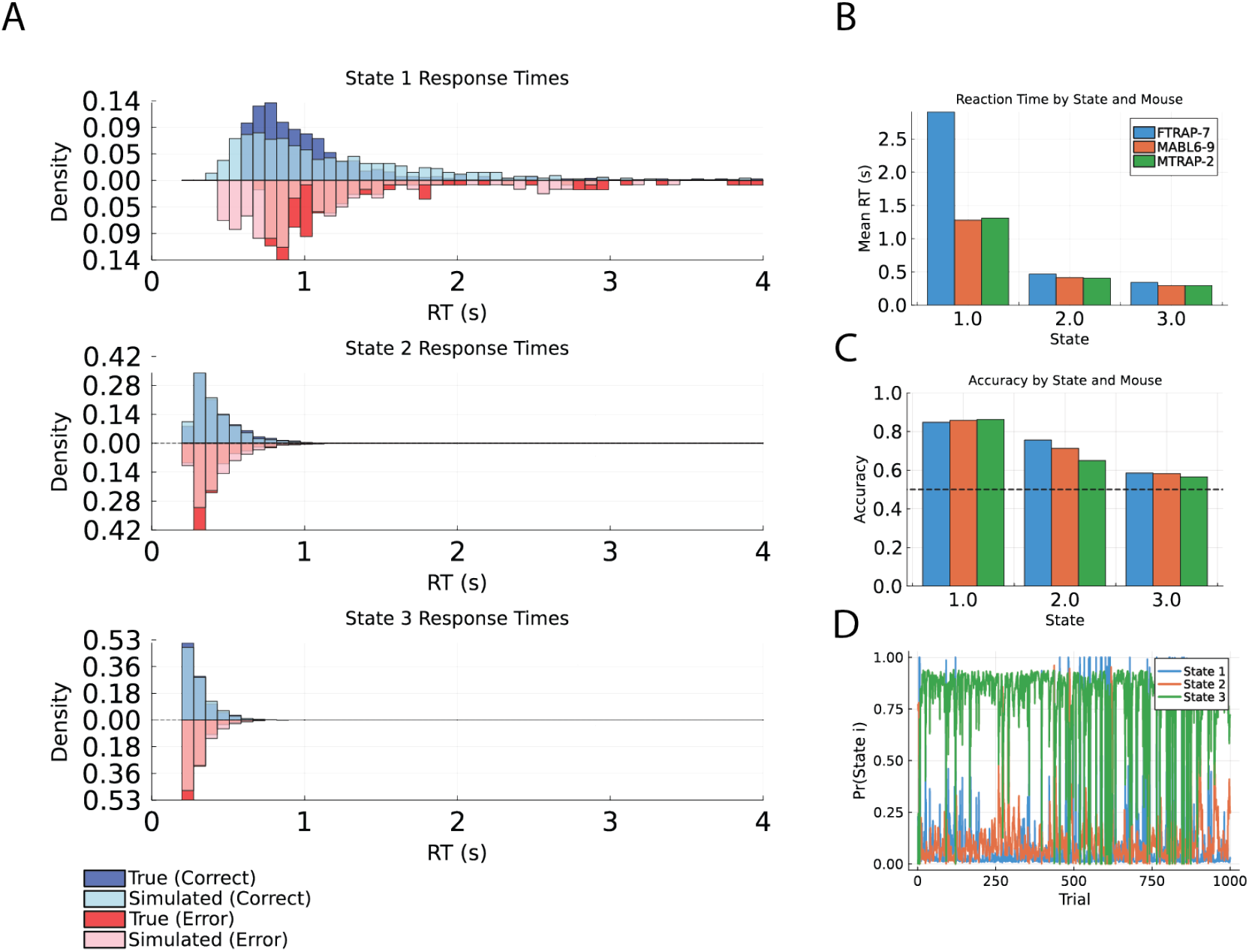
Post-hoc analysis of the HMM-DDM. **A.** Posterior predictive checks of the three learned states. The three RT distributions show distinct features, such as differences in mean and variance. **B**. Mean RT across the three states for each mouse. **C**. Mean accuracy across the three states for each mouse. **D**. Example latent state sequence posterior of the three state probabilities across trials.

State one was characterized by long RTs and high accuracy (approximately 80%), suggesting a deliberative, accuracy-oriented mode. State two reflected intermediate behavior, with moderate RTs and accuracy around 75%. In contrast, state three displayed very short RTs and low accuracy—only slightly above chance (approximately 55%)—consistent with a fast but error-prone decision-making mode. These results indicate that the latent states of the HMM-DDM capture meaningful variability in behavioral strategy, ranging from cautious and accurate to fast and impulsive responding.

Finally, we note that the model predicts that animals spend the most time in the highly inaccurate state, which aligns with the animals relatively low accuracy (approx 65%) (Figure 6 D). Interestingly, the model predicts with high confidence the animals enter these higher performing regimes corresponding to states one and two throughout the sessions, and so suggests that animals stochastically move out of a poor-performance state into states characterized by longer RTs and better accuracy.

## 8 Discussion

We introduce a state-dependent DDM, the HMM-DDM, that identifies discrete internal states associated with distinct decision policies. Our model reveals that mice dynamically transition between impulsive states (characterized by fast, low-accuracy responses) and deliberative states. These states occupy different positions along the speed–accuracy trade-off suggesting that mice may actively explore the speed–accuracy continuum. While similar phenomena have been observed based on high-resolution video tracking during perceptual decision making [12], our approach extracts this structure using substantially less behavioral data.

In addition to its methodological advantages, our framework uncovers rare states in which mice exhibit deliberation times and accuracies approaching those observed in human decision-making [19], raising new questions about the cognitive capabilities of rodent models.

These findings open several avenues for future research. A key direction is to identify the neural mechanisms underlying transitions between decision states. Recording neural activity during identified behavioral states could reveal the circuits that regulate decision policies. Furthermore, this approach may be extended to investigate how environmental variables—such as reward rate, uncertainty, or satiety—modulate state occupancy. Such extensions could help bridge computational models of decision-making with their neurobiological substrates.

There are several immediate extensions of our approach that are likely to extend the utility of the model and provide further scientific insight into decision-making. Within-trial accumulation dynamics can be modeled with impressive temporal detail [20], suggesting that we could extend our DDM observation model, for example to consider inhomogeneous temporal accumulation, to offer further insight into decision-making strategy for individual trials. Existing methods adapt powerful hierarchical linear dynamical systems models to perform within-trial inference and learning in such models [21], and we are actively integrating this approach into the framework presented here. Neural activity has also been shown to reflect within-trial accumulation dynamics [22]; although we considered behavioral measurements as our observation, using neural [23] or keypoint data [12] is likely to provide further insight into the accumulation process. Finally, we have exclusively considered a set of discrete states for the underlying latent state sequence, but a continuous state sequence, as could be modeled using a linear [24] or nonlinear dynamical system [25] would add further expressivity to our framework, yielding greater insight into the behavioral strategy switching exhibited by animals and humans.

## 9 Limitations

Although our DDM observation model is flexible and well-parameterized, it can struggle to account for especially non-normative and idiosyncratic behaviors often exhibited by certain species, especially mice. For example, mice occasionally make rapid, stimulus-independent choices that do not appear to reflect integration of sensory evidence. These responses often result in unusually fast reaction times with near-chance accuracy, which the model may misinterpret as low decision bounds or high internal noise. However, such trials may instead reflect momentary lapses in task engagement or non-decision-related motor biases.

To overcome this, we were required to systemically remove trials that met the following criteria: exceptionally low RTs and an asymmetry in the correct vs. error trial histograms, i.e., more errors than correct choices. Our experiments indicate that our approach groups these ‘outlier’ trials into a single state of behavioral strategies defined by trials poorly described by a DDM but that collectively are not very similar. Extensions of the model that can systematically address model mismatch between the DDM and specific behavioral strategies will mitigate these inefficiencies. Furthermore, in our experiments, the non-decision time variable *τ* can cause difficulties during optimization. Specifically, this variable causes a non-smooth transition in the likelihood function (any RTs below the chosen value of *τ* are assigned zero likelihood). To overcome this, we chose to enforce the minimum likelihood value to be a small positive value (*ϵ* = 1*e* − 12), which mostly solved the issue, but it is likely that more elegant fixes are available. Lastly, latent variable models by their construction are degenerate; without a ground-truth knowledge of the latent variables, it is difficult to know if a global, uniquely-identifiable, mode has been achieved. Thus, clever initialization strategies and optimization tricks should likely be employed to yield better performance.

